# Dimension-Selective Attention and Dimensional Salience Modulate Cortical Tracking of Acoustic Dimensions

**DOI:** 10.1101/2021.05.10.443391

**Authors:** Ashley E. Symons, Fred Dick, Adam T. Tierney

## Abstract

Some theories of auditory categorization suggest that auditory dimensions that are strongly diagnostic for particular categories - for instance voice onset time or fundamental frequency in the case of some spoken consonants - attract attention. However, prior cognitive neuroscience research on auditory selective attention has largely focused on attention to simple auditory objects or streams, and so little is known about the neural mechanisms that underpin dimension-selective attention, or how the relative salience of variations along these dimensions might modulate neural signatures of attention. Here we investigate whether dimensional salience and dimension-selective attention modulate cortical tracking of acoustic dimensions. In two experiments, participants listened to tone sequences varying in pitch and spectral peak frequency; these two dimensions changed at systematically different rates. Inter-trial phase coherence (ITPC) and EEG signal amplitude at the rates of pitch and spectral change allowed us to measure cortical tracking of these dimensions. In Experiment 1, tone sequences varied in the size of the pitch intervals, while the size of spectral peak intervals remained constant. Neural entrainment to pitch changes was greater for sequences with larger compared to smaller pitch intervals, with no difference in entrainment to the spectral dimension. In Experiment 2, participants selectively attended to either the pitch or spectral dimension. Neural entrainment was stronger in response to the attended compared to unattended dimension for both pitch and spectral dimensions. These findings demonstrate that bottom-up and top-down attentional mechanisms enhance the cortical tracking of different acoustic dimensions within a single sound stream.

## Introduction

Speech perception requires the mapping of continuous acoustic dimensions onto discrete linguistic categories. A central issue in speech perception is how listeners dynamically integrate and weight information from different acoustic dimensions during categorization. Prior work has shown that listeners weight acoustic dimensions according to the reliability with which each dimension distinguishes between categories (Holt et al., 2018; Toscano & McMurray, 2010). When the reliability of an acoustic dimension changes due to noise (Winn et al., 2013) or short-term changes in cue distribution (Idemaru & Holt, 2011, 2014), listeners dynamically reweight acoustic dimensions accordingly. Listeners show stable individual differences in dimensional weighting strategies (Idemaru et al., 2012; Kim et al., 2018; Kong & Edwards, 2016), which may reflect differences in auditory perceptual ability (Jasmin et al., 2019, 2020) and prior language experience (Jasmin et al., 2021). Despite the central role of dimensional weighting in speech perception, surprisingly little is known about the neural mechanisms underlying this process.

One process that may contribute to the flexibility of dimensional weighting strategies across different contexts as well as the variability between individuals is dimension-selective attention. According to some theoretical accounts of speech perception, listeners dynamically allocate attentional resources towards dimensions that are informative, and away from those that are less informative (Francis et al., 2000; Francis & Nusbaum, 2002; Gordon et al., 1993; Heald & Nusbaum, 2014; Holt et al., 2018). Thus, dimension-selective attention has been suggested to play a potential role in dimensional weighting in speech perception. Long-term prior experience with speech and language may change the salience of different acoustic dimensions, potentially accounting for the differences in dimensional weighting strategies across speakers of different languages (Jasmin et al., 2021). Although there is little direct evidence for the involvement of attention in auditory category learning, eye-tracking studies have provided extensive evidence that the salience of dimensions changes during visual category learning, with increased salience for dimensions that distinguish between categories (Blair, Watson, & Meier, 2009; Blair, Watson, Walshe, et al., 2009; Carvalho & Goldstone, 2017; Chen et al., 2013; Kim & Rehder, 2011; Rehder & Hoffman, 2005a, 2005b; Zaki & Salmi, 2019). Thus, there is robust evidence that dimension-selective attention plays a crucial role in visual category learning. However, despite the central role that dimension-selective attention plays in many theories of speech perception, there is little direct evidence that auditory dimension-selective attention exists, let alone what neural mechanisms might subserve it.

How might auditory dimension-selective attention be carried out in the brain? One possibility is that neural activity tracks variations in different acoustic dimensions, with attention enhancing representations of attended dimensions and potentially suppressing representations of unattended dimensions. Attention-driven enhancement of neural tracking has been demonstrated in studies of object-based auditory selective attention. For instance, prior work has shown that neural activity synchronizes with low-frequency fluctuations in the amplitude (Ding et al., 2016; Horton et al., 2013; Luo & Poeppel, 2007; Obleser & Kayser, 2019; Zoefel et al., 2018) and pitch contour (Teoh et al., 2019) of continuous speech. Attention has been shown to enhance this cortical tracking for attended versus ignored speech streams (Ding et al., 2016; Kerlin et al., 2010; Reetzke et al., 2021; Zion Golumbic et al., 2013). Moreover, studies using non-verbal stimuli have shown that attention strengthens neural entrainment to target relative to distractor tone sequences (Elhilali et al., 2005; Laffere et al., 2020, 2021). Overall, these studies suggest that attention can modulate the extent to which the auditory system tracks variations in different sound streams. Here we suggest that a similar mechanism could underlie attention to acoustic dimensions within a single sound stream.

There is some initial evidence that attention may modulate the cortical tracking of different acoustic dimensions. For example, Costa-Faidella et al., (2017) had participants listen to single auditory streams that varied in duration and intensity at different rates; they either responded as to whether five consecutive tones were long or short (duration task) or silently counted the number of loud tones (intensity task). Neutral entrainment was stronger for the attended compared to the ignored dimension. However, the dimension to which attention was directed was not independent from a difference in task across conditions, and so the condition effect on neural tracking could have partially reflected task demands. Moreover, in the absence of a passive listening condition, it is not possible to discern whether this finding reflects neural enhancement of the attended dimension, or suppression of the unattended dimension. Therefore, it is still unclear whether the listeners are increasing the gain on the attended dimension or actively inhibiting task-irrelevant information in unattended dimension. Disentangling the effects of enhancement versus suppression will provide crucial insight into the mechanisms underpinning auditory dimension-selective attention.

In addition to being modulated by top-down attention, cortical tracking of acoustic dimensions may also be enhanced by bottom-up attentional salience. Multiple acoustic dimensions contribute to the perceptual salience of natural sounds (Huang & Elhilali, 2017; Zhao et al., 2019). Salient changes in one or more of these acoustic dimensions can modulate physiological measures of attentional orienting such as skin conductance response (Siddle et al., 1984) and pupil dilation response (Bala & Takahashi, 2000; Liao et al., 2016; Marois et al., 2018; Wetzel et al., 2016; but see Zhao et al., 2019), with the magnitude of the response varying in proportion to the size of the change (Marois et al., 2018; Wetzel et al., 2016). Prior EEG work has also shown that both the mismatch negativity (MMN) and P3 responses, associated with detection of acoustic change and orientation of attention respectively, are sensitive to the magnitude of the change along multiple acoustic dimensions (Escera et al., 1998; Berti et al., 2004; Rinne et al., 2006; Schröger, 1996). Salient acoustic changes can also influence the degree of neural entrainment to acoustic streams, with decreased entrainment to attended streams following salient background sounds (Huang & Elhilali, 2020) and increased entrainment to acoustic melodies following deviations in pitch, timbre and intensity (Kaya et al., 2020). These studies suggest that salient changes in a sound stream along a number of different dimensions can attract attention to the sound stream. However, it remains unclear whether dimensional salience can attract attention to specific acoustic dimensions within a sound stream, as revealed by enhanced neural entrainment to increased salience along a dimension.

To investigate whether cortical tracking of acoustic dimensions is modulated by dimensional salience (Experiment 1) and dimension-selective attention (Experiment 2), we used a frequency tagging approach (e.g., Ding et al., 2016; Nozaradan et al., 2011), where changes in different dimensions are tagged to fixed rates. In each experiment, listeners heard sequences of synthesized complex tones that varied in pitch (fundamental frequency) and in spectral peak frequency, each at a given fixed rate. In Experiment 1, the salience of the pitch dimension was manipulated by altering the pitch step sizes (1 versus 2 semitones) between blocks, while the step sizes of the spectral peak frequency remained constant (2 semitones). Based on previous research showing that responses to pitch deviants vary as a function of step size (Berti et al., 2004; Marois et al., 2018; Wetzel et al., 2016), we hypothesized that neural entrainment would be modulated by increased salience along the pitch dimension. Thus, we predicted that stronger inter-trial phase-locking at the rate tagged to pitch changes would be observed in blocks with larger pitch step sizes (2 semitones) compared to smaller pitch step sizes (1 semitone).

In Experiment 2, the tone sequences were identical across conditions, while only the focus of attention varied. In two attention conditions, listeners either attended to variations in the pitch, or the spectral peak frequency dimension, while ignoring variations in the other dimension. In a third ‘neutral’ condition, listeners monitored for occasional quiet tones. This condition allowed us to test whether the effects of attention were due to attentional enhancement of the attended dimension, or suppression of the unattended dimension. We hypothesized that dimension-selective attention would result in enhanced neural entrainment to a dimension when it was attended, and potentially suppressed neural entrainment to a dimension when it was unattended.

## Experiment

### Methods

#### Participants

Twenty-nine participants (14 female, 15 male) between the ages of 19-59 with no known hearing impairments and no diagnosis of a language or learning disorder took part in this experiment. This sample size was powered to detect a medium effect (d = 0.5) between 2-semitone versus 1-semitone pitch step size conditions (α = 0.05, β = 0.8).

Three participants were subsequently excluded. Two of the three were excluded as a result of insufficient EEG data, one due to excessive EEG artefacts resulting in the loss of over 50% of trials in multiple conditions and another due to a technical error that resulted in the loss of data from one condition. The third participant was excluded on the basis of poor behavioral performance (< 10% hit rate and >100 false alarms).

The final sample consisted of 26 participants (12 female, 14 male) with a mean age of 31.65 years (standard deviation = 10.06 years). Native languages spoken by participants included English (19), Polish (2), Greek (1), Japanese (1), Mandarin (1), Romanian (1), and Russian/Uzbek (1). Twelve participants reported receiving some form of musical training, ranging from 3-20 years (mean = 10.96, standard deviation = 5.60).

The Ethics Committee in the Department of Psychological Sciences at Birkbeck, University of London approved all experimental procedures. Informed consent was obtained from all participants. Participants were compensated for their participation in the form of course credits, or payment at a standard rate.

#### Design

In this experiment, participants listened to isochronous sequences of complex tones that varied in pitch and spectral peak frequency at different rates (see Figure 1). The rates at which the pitch and spectral peak changed were counterbalanced within participants to ensure that any effect of dimensional salience was not due to differences in neural entrainment to different presentation rates. Pitch salience was manipulated by varying the pitch step sizes. This resulted in an experimental design with two pitch step size conditions (1 versus 2 semitones) and two varying acoustic dimensions (pitch versus spectral peak).

**Figure 1.**
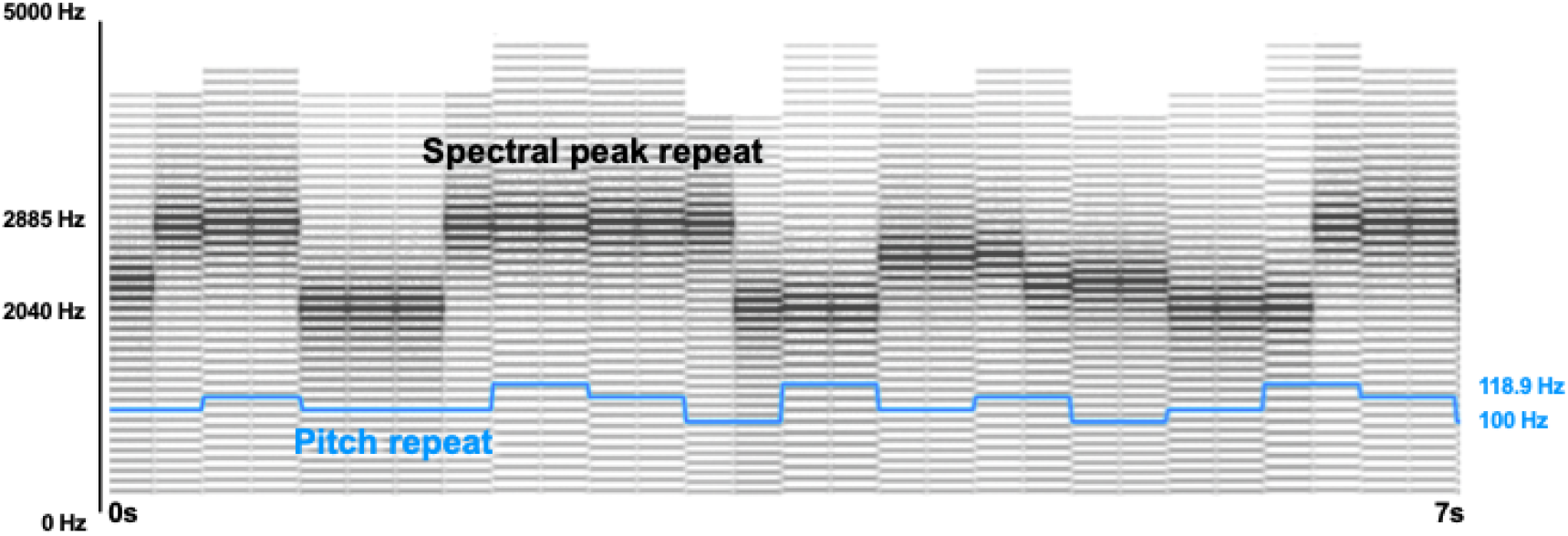
Spectrogram showing a cropped portion of a stimulus varying in pitch (blue line) and spectral peak. In this example, pitch step sizes are separated by 1 semitone (min = 100 Hz, max = 118.9 Hz) and spectral peak step sizes are separated by 2 semitones (min = 2040 Hz, max = 2885 Hz).

#### Stimuli

The stimuli consisted of 250 ms complex tones (40 harmonics and a 15-ms linear on/off ramp) with a single spectral peak (Smith, 2007). Two 2-dimensional stimulus spaces were created. In one, complex tones with one of four fundamental frequencies (each separated by two semitones, 91.13 Hz, 102.29 Hz, 114.82 Hz, 128.87 Hz) were modulated by one of four spectral peaks (again separated by two semitones, 2040.00 Hz, 2289.82 Hz, 2570.24 Hz, 2884.99 Hz). The second stimulus space only differed from the first in that the fundamental frequencies were separated by a single semitone (100 Hz, 105.94 Hz, 112.25 Hz, 118.92 Hz). Importantly, although the pitch step sizes varied, the mean fundamental frequency of the two spaces was the same (109.28 Hz).

These two sets of tones were concatenated without pause to form 120-second sequences (480 tones). Crucially, pitch and spectral peak changed at different rates (1.33 Hz and 2 Hz). When pitch varied at 2 Hz, spectral peak varied at 1.33 Hz. When pitch varied at 1.33 Hz, spectral peak varied at 2 Hz (Figure 1). The order of the tones within each sequence was pseudorandomized such that tones had to change in pitch and spectral peak at the specified rate, with the exception of 20 repeated segments that occurred within each dimension. These repetitions were instances in which the dimension did not change at the expected rate. An example of a repetition would be if pitch was typically changing every 2 tones (2Hz), but at the repetition did not change until after 4 tones. Repetitions were randomly inserted into the sequence, with the condition that two repetitions could not occur consecutively. The repetitions were included for comparability with Experiment 2, and are not directly relevant to Experiment 1; Experiment 1 participants were not alerted to their presence, and the repetitions were task-irrelevant.

The participants’ explicit behavioral task was to respond to quiet ‘oddball’ tones, where the amplitude of 3-4 randomly selected tones was decreased by 25% (−12.0 dB, see below). Oddball timing was randomized in each sequence, with the exception that oddballs could not occur in the first or last 1.5 seconds of the sequence, and could not occur consecutively. The same sequences were presented to all participants, but with the order of conditions counterbalanced across participants.

This resulted in four conditions, each consisting of four sequences that varied in a) pitch step size (1 semitone or 2 semitone difference between pitch steps) and b) dimension change rate (pitch at 1.33 Hz/spectral peak at 2 Hz, or pitch at 2 Hz/spectral peak at 1.33 Hz). Stimulus presentation was blocked by condition, but with the order of conditions counterbalanced across participants.

Stimuli were presented using PsychoPy3 (v 3.2.3) and the sound delivered via ER-3A insert earphones (Etymotic Research, Elk Grove Village, IL) at a level of 80 dB SPL.

#### Procedure

Prior to the EEG task, participants completed a short practice task outside of the EEG booth. Here, they listened to two short sequences (48 seconds), each consisting of two oddball tones, and pressed the keyboard space bar when they detected those tones. Participants received visual feedback on their performance. If they failed to detect at least two of the four quiet oddball tones, or if they had more than four false alarms on the second sequence, the task was explained to them a second time and they were asked to complete a second practice block on a different set of sequences. All participants were able to pass the practice task.

In the main EEG task, participants listened to each 120-second sequence and responded via a keyboard press when they detected the oddball tones. This task was chosen in order to keep participants awake and alert throughout the task without focusing attention on one of the two dimensions of interest. In contrast to the training, participants did not receive visual feedback on their responses in this task.

#### EEG Data Acquisition and Preprocessing

EEG data was recorded from 32 Ag-Cl active electrodes using a Biosemi™ ActiveTwo system with electrodes positioned according to the 10/20 montage. Data were recorded at a sampling rate of 16,384 Hz and digitized with 24-bit resolution. Two external reference electrodes were placed on the earlobes for off-line re-referencing. Impedance was kept below 20 kΩ. Triggers marking the beginning of each trial (every 6 tones, or 1.5 seconds) were recorded from trigger pulses sent to the data collection computer.

The data were down sampled to 512 Hz and re-referenced to the average of the earlobe reference electrodes. A low-pass zero-phase sixth-order Butterworth filter with a cutoff of 30 Hz was applied. A high-pass fourth-order zero-phase Butterworth filter with a cut-off of 0.5 Hz was then applied and the data epoched (1.5-seconds for analysis of phase and 30-seconds for analysis of signal amplitude) based on the recorded trigger pulses. Independent component analysis (ICA) was conducted to correct for eye blinks and horizontal eye movements. Components corresponding to eye blinks and movements were identified and removed based on visual inspection of the time courses and topographies.

Two measures of neural entrainment were computed: inter-trial phase coherence (ITPC) and signal amplitude. Prior to calculation of ITPC, any remaining artifacts exceeding +/- 100 μV were rejected. A Hanning-windowed fast Fourier transform was then applied to each 1.5-second epoch. The complex vector at each frequency was converted to a unit vector and averaged. The length of the average vector provides a measure of ITPC, which ranges from 0 (no phase consistency) to 1 (perfect phase consistency).

For analysis of signal amplitude, the 30-second epochs were averaged for each condition and transformed into the frequency domain using a fast Fourier transform. The resulting frequency spectrum represents the amplitude (in microvolts) at each frequency, with a frequency resolution of 1/30 (0.033 Hz). The frequency spectrum was then normalized by taking the difference between the amplitude at each frequency and the mean amplitude of the four neighboring frequencies (Nozaradan et al., 2011) in order to reduce the contribution of noise and other ongoing neural activity from the EEG signal.

All EEG data processing and analysis were carried out in Matlab (MathWorks, Inc) using the FieldTrip M/EEG analysis toolbox (Oostenveld et al., 2011) in combination with in-house scripts.

### Data Analysis

#### Behavioral Data

The primary purpose of the behavioral task was to keep participants alert throughout the presentation of the stimuli, without having them explicitly attend to either of the two dimensions. Therefore, the task was designed to be easy, and performance was expected to be near ceiling. Nonetheless, the proportion of hits and false alarms was calculated for the 2-semitone and 1-semitone pitch step size conditions. The proportion of false alarms was defined as the number of responses that occurred outside of 1.25 seconds following an oddball, divided by the total number of non-oddball tones occurring outside of the target time windows. The proportion of hits and false alarms was converted to d-prime for analysis, with the loglinear approach used to avoid infinite scores (Hautus, 1995). Wilcoxon signed-rank tests were used to test for an effect of pitch step size on performance.

#### EEG Data

For both phase and amplitude measures, we extracted data from the 9 channels with the maximum signal when averaged across all 4 conditions, the 2 frequencies (1.33 Hz and 2 Hz) relevant for assessment of neural tracking of stimulus dimensions, and all 26 participants. (Note that this choice of channels, which was based on collapsing across conditions, was orthogonal to our analysis, which was a comparison between conditions.) This resulted in the same set of frontocentral channels (AF3, AF4, F3, F4, Fz, FC1, FC2, Cz, C3) for both phase and amplitude measures. The data were averaged across these channels prior to statistical analysis. We then collapsed the data across the two different rates of dimension change to determine the degree of neural entrainment to pitch and spectral peak dimensions. Wilcoxon signed-rank tests were used to compare the effect of pitch step size on neural entrainment to pitch and spectral peak. False Discovery Rate was used to correct for multiple comparisons using the Benjamini and Hochberg procedure (Benjamini & Hochberg, 1995). Processed data are available at: osf.io/c5s6u/.

### Results

#### Behavioral

As expected, performance was near ceiling in both the small pitch step size (median d-prime = 4.863) and large pitch step size (median d-prime = 4.599) conditions. The difference between the two conditions was statistically significant (V = 43, z = −252 p = 0.012). The difference in behavioral performance was unexpected but could suggest that larger pitch step sizes are more distracting, resulting in worse performance. However, upon closer investigation, this difference may also be due to the different proximity of the quiet oddball tones in the different conditions. Specifically, we noticed that there were 3 instances where oddball tones occurred within the same 1.25 second window as another oddball tone. Two of these occurred in the 2-semitone pitch step size condition and one occurred in the 1-semitone pitch step size condition. Removing responses to both oddball tones within these time windows along with two extra randomly-selected responses to oddball tones in the 1-semitone condition (to ensure that the total proportion of hits and false alarms was balanced across the two conditions) from all participants resulted in no significant difference in behavioral performance between small (median d-prime = 4.89) and large (median d-prime = 4.80) pitch step size conditions (V = 44, z = −1.54, p = 0.129).

#### EEG

Figure 2 displays neural entrainment to pitch and spectral dimensions at 1-semitone and 2-semitone pitch step sizes for ITPC (A) and signal amplitude (B). For both measures, neural entrainment to pitch changes was stronger for the 2-semitone compared to 1-semitone condition (ITPC: V = 284, z = 2.760, p_(corrected)_ = 0.009, amplitude: V = 300, z = 3.160, p_(corrected)_ = 0.004). As predicted, neural entrainment to spectral peak changes presented at the same rates did not differ as a function of pitch condition (ITPC: V = 121, z = −1.38, p_(corrected)_ = 0.231, amplitude: V = 148, z = −0.698, p_(corrected)_ = 0.499). In fact, there was no significant difference in neural entrainment to pitch versus spectral dimension in the 1-semitone condition (ITPC: V = 186, z = 0.267, p_(corrected)_ = 0.803; amplitude: V = 199, z = 0.597, p_(corrected)_ = 0.754), while in the 2-semitone condition neural entrainment was stronger to the pitch compared to spectral dimension (ITPC: V = 319, z = 364, p_(corrected)_ < 0.001; amplitude: V = 326, z = 3.820, p_(corrected)_ < 0.001).

**Figure 2.**
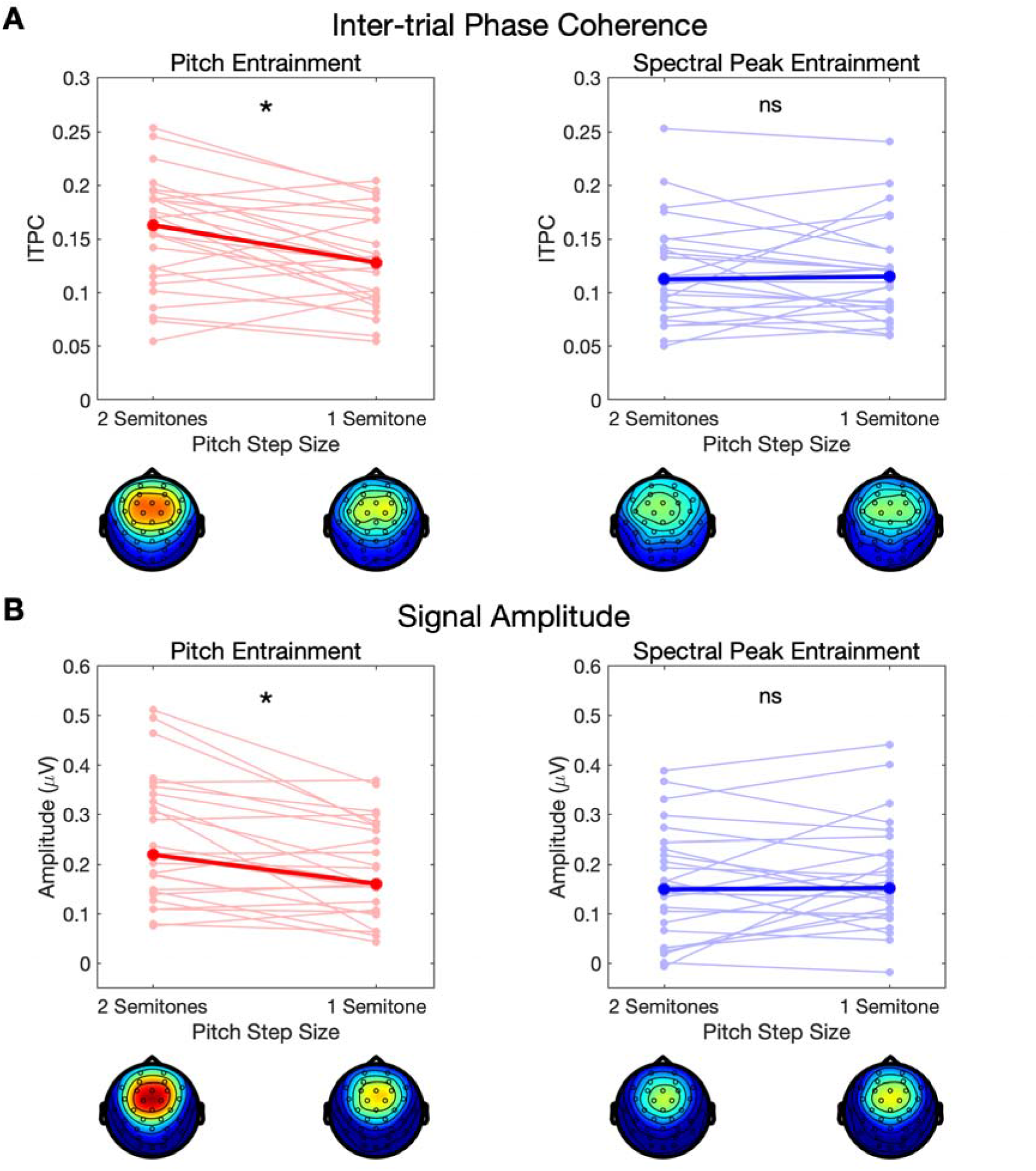
(A) Inter-trial phase coherence in response to variations in the pitch (red) and spectral peak (blue) dimensions for each pitch step size (x-axis). The values displayed represent the average across the frontocentral channels selected for analysis. Individual participants are represented in light red/blue lines and the median value represented by the dark red/blue lines, with topographical plots (scale: 0.05 – 0.2) for each condition displayed below the x-axis. (B) Amplitude at the frequencies corresponding to variations in the pitch (left) and spectral peak (right) dimensions for each pitch step size (x-axis) across the frontocentral channels selected for analysis. Individual participant data are represented by light red/blue lines and the median is represented by the dark red/blue lines, with topographical plots (scale = 0.05 – 0.3) below the x axis. For both measures, neural entrainment to the pitch dimension was stronger for the 2 compared to 1 semitone pitch step size condition. No effect of pitch step size was observed on neural entrainment to the spectral peak dimension.

## Experiment 2

### Methods

#### Participants

Twenty-seven participants between the ages of 17-52 took part in the behavioral training tasks. This sample size was sufficient to provide 80% power to detect a medium effect size (d = 0.5) between conditions. Of these participants, 6 failed to reach the performance threshold required to pass the training. The final sample consisted of 21 participants (7 female, 14 male) between the ages of 17-52 (mean age = 34.71, standard deviation = 8.71) with no known hearing impairments and no language or learning disorders. Native languages spoken by participants included English (12), Italian (4), Arabic (1), Croatian (1), German (1), Hungarian (1), and Portuguese (1). Due to the challenging nature of the task, we primarily recruited individuals with musical training. Nineteen participants reported receiving some form of musical training, with the number of years of musical training ranging between 4-20 years (mean = 13.48, standard deviation = 4.87). This final sample size was sufficient to supply 71.5% power to detect a medium effect size.

The Ethics Committee in the Department of Psychological Sciences at Birkbeck, University of London approved all experimental procedures. Informed consent was obtained from all participants. Participants were compensated for their participation in the form of course credits or payment at a standard rate.

#### Design

This experiment consisted of two phases: behavioral training and EEG testing. The behavioral training consisted of two tasks. The single dimension training task involved attending to variations in one dimension while the other dimension remained constant. The dimension-selective attention training task involved attending to one dimension while ignoring variations in the other dimension. In the EEG testing phase, participants completed a longer version of the dimension-selective attention task while EEG was recorded. In each task, there were three attention conditions: attend pitch, attend spectral peak, and a ‘neutral’ attention condition where participants performed the amplitude oddball detection task of Experiment 1, e.g., detecting occasional oddball tones that were reduced in amplitude relative to the other tones in the sequence. The rate of pitch and spectral peak change was counterbalanced within participants to ensure that any effect of attention was not due to differences in neural entrainment to different presentation rates. This resulted in an experimental design with three attention conditions (attend pitch, attend spectral peak, neutral) and, for neural measures, two phase-locked dimensions (pitch and spectral peak).

#### Stimuli

The results from Experiment 1 suggested that the salience of the pitch and spectral peak dimensions was relatively balanced, on average, in the 1-semitone pitch step size condition. Therefore, in this experiment, we used pitch step sizes of 1 semitone (100 Hz, 105.94 Hz, 112.25 Hz, 118.92 Hz) while the spectral peak step sizes varied by 2 semitones (2040.00 Hz, 2289.82 Hz, 2570.24 Hz, 2884.99 Hz).

#### Training Stimuli

For the single dimension training, we created 48-second sequences in which only one dimension varied while the other dimension remained constant. In the attend pitch condition, the spectral peak was constant at 2040 Hz while F0 varied at either 1.33 Hz or 2 Hz. In the attend spectral peak condition, F0 was constant at 100 Hz while only the spectral peak varied every two (2 Hz) or three (1.33 Hz) tones. Eight repetitions, or instances in which the attended dimension did not change at the expected rate, were inserted into each sequence. In the ‘neutral’ attention condition, F0 was held constant at 100 Hz and spectral peak at 2040 Hz. In this condition only, two randomly selected tones were reduced in amplitude by −12 dB. This resulted in five conditions: attend pitch (pitch varying at 1.33 Hz), attend pitch (pitch varying at 2 Hz), attend spectral peak (spectral peak varying at 1.33 Hz), attend spectral peak (spectral peak varying at 2 Hz), and neutral attention (amplitude oddball detection). Five tone sequences were generated for each condition.

For the dimension-selective attention training, 48-second sequences were created in which the two dimensions varied at different rates. In half of the sequences, pitch varied at 2 Hz and spectral peak varied at 1.33 Hz; in the other half of the sequences, spectral peak varied at 2 Hz and pitch at 1.33 Hz. Eight repetitions along each dimension were inserted into each training sequence. In addition, 3-4 tones were decreased in amplitude by −12 dB in all sequences. The tones that were decreased in amplitude were randomly selected, with the exception that they could not occur during a repetition, or at the tone immediately preceding or following a repetition. Six conditions were created by crossing attention condition (pitch, spectral peak, neutral) with dimension change rate (pitch and spectral peak): attend pitch (pitch at 1.33 Hz/spectral peak at 2 Hz), attend pitch (pitch at 2 Hz/spectral peak at 1.33 Hz), attend spectral (pitch at 1.33 Hz/spectral peak at 2 Hz), attend spectral (pitch at 2 Hz/spectral peak at 1.33 Hz), neutral (pitch at 1.33 Hz/spectral peak at 2 Hz), and neutral (pitch at 2 Hz, spectral peak at 1.33 Hz). Five tone sequences were generated for each condition.

#### EEG Stimuli

The characteristics of the stimuli for the main EEG task were identical to those of the small pitch step size condition of Experiment 1. As with Experiment 1, the 250 ms tones were concatenated into 120 s sequences with pitch and spectral peak dimensions varying at different rates (1.33 Hz or 2 Hz). Twenty repetitions (instances in which the dimension did not change at the expected rate) were inserted into each sequence; these were task-relevant for the pitch and spectral attention conditions. Additionally, 3-4 randomly selected tones were decreased in volume by −12 dB; as previously, these were task-relevant in the neutral conditions. The locations of the quiet oddball tones were pseudorandomly selected, with the exceptions that the oddball tones could not occur in the first or last 1.5 seconds of each sequence, nor could they occur during or immediately before or after a repetition. For this experiment we also increased the minimum temporal interval between successive oddball tones such that there needed to be at least 6 standard-amplitude tones between successive oddball tones.

As with the dimension-selective training stimuli, six conditions were created by crossing attention condition (pitch, spectral peak, neutral) with dimension change rate (pitch and spectral peak): attend pitch (pitch at 1.33 Hz/spectral peak at 2 Hz), attend pitch (pitch at 2 Hz/spectral peak at 1.33 Hz), attend spectral (pitch at 1.33 Hz/spectral peak at 2 Hz), and attend spectral (pitch at 2 Hz/spectral peak at 1.33 Hz), neutral (pitch at 1.33 Hz/spectral peak at 2 Hz), and neutral (pitch at 2 Hz, spectral peak at 1.33 Hz). Each of the six conditions consisted of four sequences of tones. Stimulus presentation was blocked by condition, with the order of conditions counterbalanced across participants.

### Procedure

#### Behavioral Training

Prior to the EEG task, participants completed two short behavioral training exercises outside of the EEG booth. In the first training exercise, participants listened to shortened sequences (48 seconds) in which only a single dimension varied. First, participants were familiarized with the two dimensions. The pitch dimension was described to participants as how ‘high or low’ the sound was. The spectral peak dimension was described to participants as ‘brightness’. Participants were provided examples of the variations in each dimension that they could listen to multiple times before progressing to the task. For attend pitch and spectral peak conditions, participants were told which dimension to attend to, and how often that dimension would change. Their task was to press a button to respond when they detected repetitions in the attended dimension. In the neutral condition, participants were instructed to listen out for and respond to occasional quiet tones. Participants received feedback on their performance. Participant’s total score was displayed on the screen, with +1 point for every correct detection and −1 for every false alarm. For the attend pitch and spectral peak conditions, if participants received a total score of < 7/8, they received another training block for that condition. If participants met or exceeded the threshold, they moved onto the next condition. For the neutral condition, if participants missed more than one quiet tone (out of two) or had more than one false alarm, they received another block of training on that condition.

In the second training exercise, participants listened to shortened sequences (48 seconds) in which *both* pitch and spectral peak dimensions changed at different rates. Quiet tones were also embedded in these training sequences. Prior to each sequence, participants were told which dimension to attend to, and how often that dimension would change. The task was the same as the single-dimension training, except that participants also had to ignore changes in the unattended dimension. Participants received visual feedback on their performance identical to that in the single dimension training. Participant’s total score was displayed on the screen, with +1 point for every correct detection and −1 for every false alarm. For the attend pitch and spectral peak conditions, if participants received a total score of < 6/8, they received another training block for that condition. If participants met or exceeded the threshold, they moved onto the next condition. For the neutral condition, if participants missed more than one quiet tone (out of three or four) or had more than one false alarm, they received a second block of training on that condition.

Due to the COVID-19 pandemic, in-lab testing was suspended part-way through the experiment. When testing could safely be resumed, the behavioral training tasks were moved from in-lab to online to minimize the amount of time the participants and researchers had to interact in the lab. Minor variations existed between the in-lab and online training, and are described in the Supplementary Materials.

#### EEG Testing

In the main EEG task, participants listened to 120-second sequences in which both pitch and spectral peak dimensions varied. There were 6 conditions (2 dimension change rates x 3 attended dimensions). Stimulus presentation was blocked by attended dimension in order to minimize switching costs, with the order counterbalanced across participants. At the start of each block, participants were instructed to detect repetitions in the attended dimension (attend pitch and spectral peak conditions) or to detect the quiet tones. In the attend pitch and attend spectral blocks only, participants were also told how often they could expect the attended dimension to vary. For example, in an attend pitch block in which pitch varied at 2 Hz, participants would hear the instructions: “In this block, your task is to attend to pitch, which will change every 2 sounds. Press the trigger button when you hear a repetition in pitch.” In contrast to the training, no visual feedback was provided. Participants made their responses by pressing the trigger on an Xbox One game controller. Stimuli were presented using PsychoPy3 (v 3.2.3) and the sound delivered via ER-3A insert earphones (Etymotic Research, Elk Grove Village, IL) at a level of 80 dB SPL. Each block lasted between 8-10 minutes. In between each sequence, participants had the option of taking a short self-paced break. The total duration of the EEG task was approximately 1 hour.

#### Data Acquisition and Analysis

EEG data acquisition and preprocessing procedures were identical to those of Experiment 1.

##### 2.1.5.1 Behavioral

In the attend pitch and attend spectral peak conditions the proportion of hits and false alarms was calculated for each attention condition. Hit rate was defined as responses occurring within 1.25 seconds following a repetition divided by the total number of repetitions. False alarm rate defined as the number of responses occurring outside of the 1.25-second target window, divided by the total number of non-repetitions (instances where the dimension changed at the expected rate). In the neutral condition, the proportion of hits and false alarms was computed in the same manner as in Experiment 1. All scores were converted to d-prime for analysis with the loglinear approach used to avoid infinite values (Hautus, 1995). Statistical analysis was conducted separately for the dimension-selective attention (attend pitch, attend spectral peak) and amplitude oddball detection tasks since behavioral performance on the two tasks was not directly comparable. A Wilcoxon signed-rank test was used to compare performance in the attend pitch and attend spectral peak conditions, with False Discovery Rate used to correct for multiple comparisons via the Benjamini and Hochberg procedure (Benjamini & Hochberg, 1995).

#### EEG

ITPC and signal amplitude were extracted over the 9 frontocentral channels identified in Experiment 1 (AF3, AF4, F3, F4, Fz, FC1, FC2, Cz, C3). The data were averaged across channels prior to statistical analysis. We then collapsed the data across the two different dimension change rates to determine the degree of neural entrainment to pitch and spectral peak dimensions in each attention condition. Wilcoxon signed-rank tests were used to compare the different attention conditions (attend pitch, attend spectral peak, neutral) on pitch and spectral peak entrainment. False Discovery Rate was used to correct for multiple comparisons using the Benjamini and Hochberg procedure (Benjamini & Hochberg, 1995). Processed data are available at: osf.io/c5s6u/.

### Results

#### Behavioral

##### Training

In the single dimension training, participants who passed the training required between 1-5 blocks of training to reach the performance threshold. Hit rate ranged from 0.63-1.0 (median = 1.0) in the attend spectral peak condition and 0.81-1.00 (median = 1.0) in the attend pitch condition. The number of false alarms ranged from 0-5 (median = 0) in the attend spectral peak condition and 0-2 (median = 0) in the attend pitch condition.^1^ In the neutral condition, hit rate was at ceiling (1.0) for all participants with between 0-1 (median = 0) false alarms. In the dimension-selective attention training, hit rate ranged from 0.75-1.0 (median = 0.94) in the attend spectral peak condition and 0.86-1.0 (median = 0.94) in the attend pitch condition. The number of false alarms in both conditions ranged from 0-3 (median = 1). In the neutral condition, hit rate was at ceiling (1.0) for all participants with between 0-1 false alarms (median = 0). No significant differences were observed between attend pitch and attend spectral peak conditions on either task (p > 0.05). These data show that participants understood the task instructions and could selectively attend to the different acoustic dimensions.

#### EEG Task

Figure 3 shows task performance (d-prime) in attend pitch and attend spectral peak conditions. There was no significant difference in performance between the two conditions (V = 162, z = 1.08, p = 0.111), suggesting that, on balance, task difficulty was matched across the two dimensions. In the neutral condition, performance was high (median d-prime = 4.857), comparable to the behavioral performance observed in Experiment 1.

**Figure 3.**
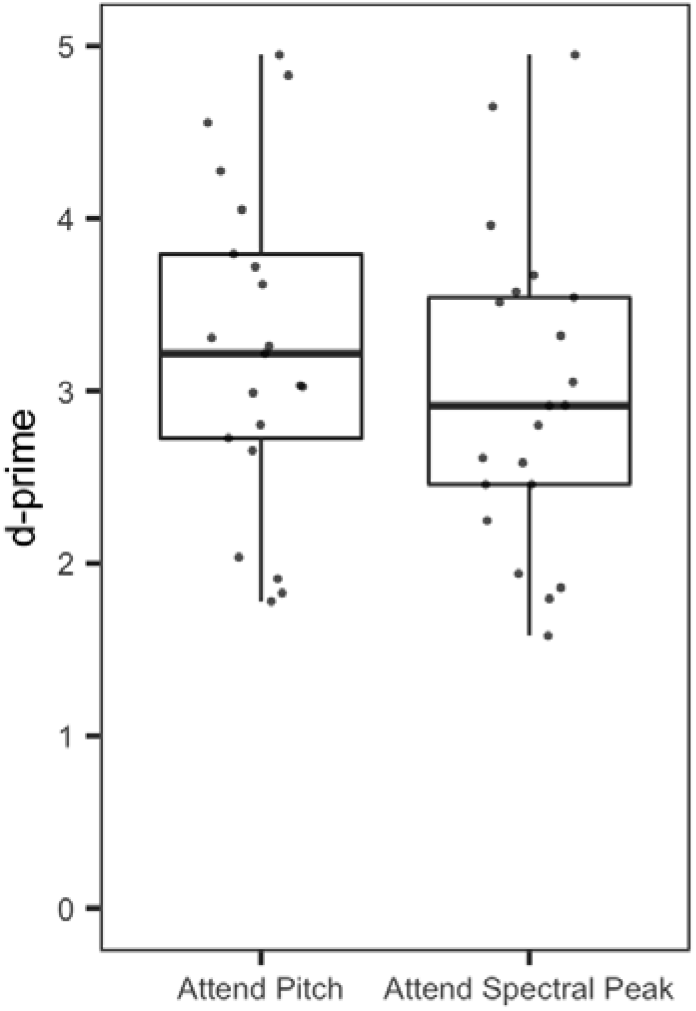
Behavioral performance (d-prime) in the attend pitch and attend spectral peak conditions. No significant differences in performance were observed between the two conditions.

#### EEG

As shown in Figure 4, neural entrainment to pitch was stronger in the attend pitch compared to the attend spectral peak condition (ITPC: V = 187, z = 2.490, p_(corrected)_ = 0.017, amplitude: V = 188, z = 2.520, p_(corrected)_ = 0.017), as well as in the attend pitch compared to neutral condition (ITPC: V = 209, z = 3.250, p_(corrected)_ = 0.002, amplitude: V = 208, z = 3.220, p_(corrected)_ = 0.002). There was also significantly greater neural entrainment to pitch in the attend spectral peak versus the neutral conditions as assessed via ITPC (V = 179, z = 2.210, p_(corrected)_ = 0.035). However, this effect was not observed for signal amplitude (V = 162, z = 1.620, p_(corrected)_ = 0.133).

**Figure 4.**
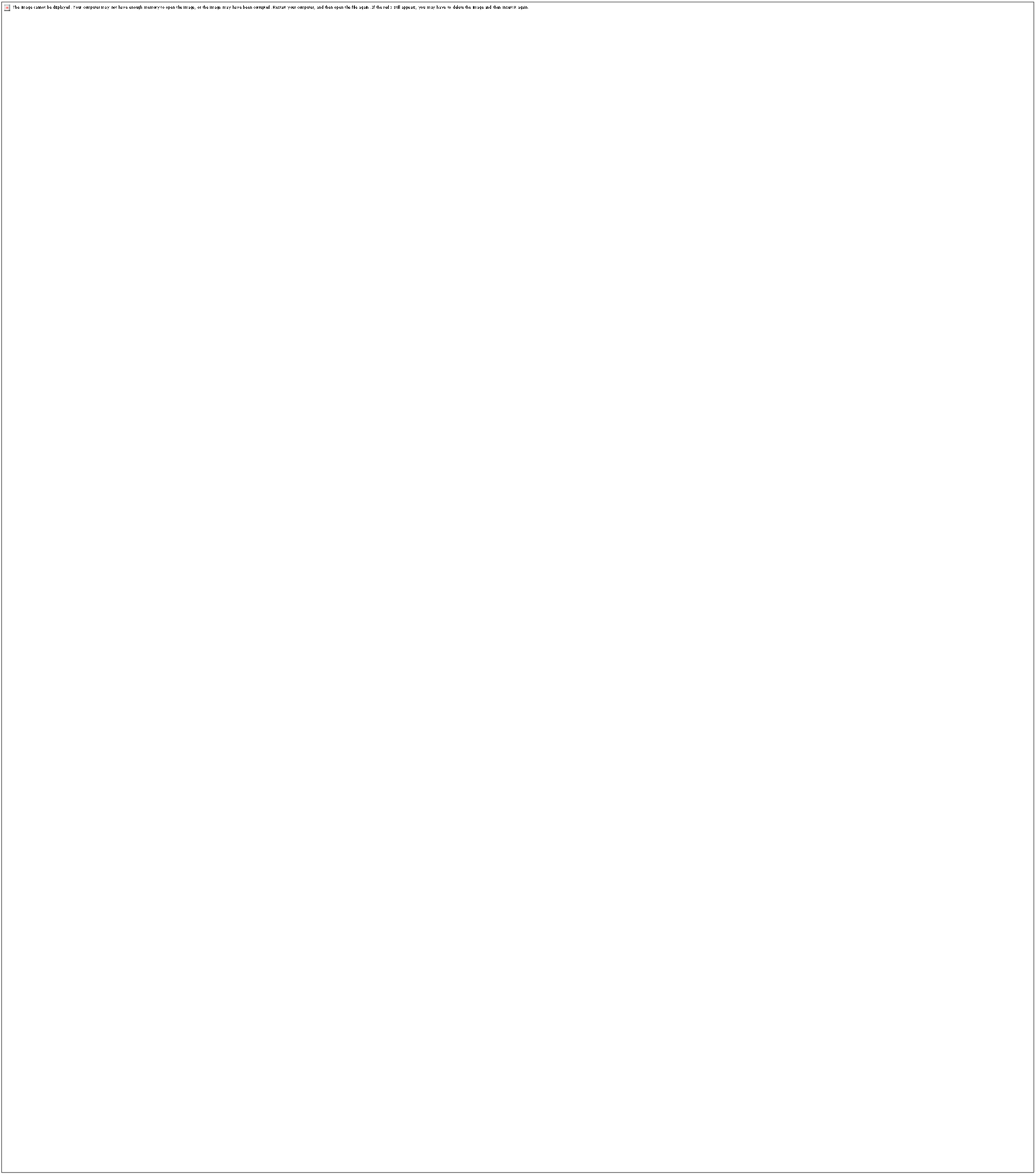
(A) Inter-trial phase coherence in response to variations in the pitch (left) and spectral peak (right) dimensions for each attention condition (x-axis). The values displayed represent the average across the frontocentral channels identified in Experiment 1. Individual participants are represented in light red/blue lines and the median value represented by the dark red/blue lines, with topographical plots (scale: 0.05 – 0.2) for each condition displayed below the x-axis. (B) Amplitude at the frequencies corresponding to variations in the pitch (left) and spectral peak (right) dimensions for each attention condition across the frontocentral channels selected for analysis. Individual participant data are represented by light red/blue lines and the median is represented by the dark red/blue lines, with topographical plots (scale = 0.05 – 0.3) below the x axis.

Neural entrainment to the spectral peak dimension was stronger in the attend spectral peak compared to the attend pitch condition (ITPC: V = 193, z = 2.690, p_(corrected)_ = 0.013, amplitude: V = 187, z = 2.490, p_(corrected)_ = 0.017), and in the attend spectral peak compared to neutral condition (ITPC: V = 209, z = 3.250, p_(corrected)_ = 0.002, amplitude: V = 199, z = 2.900, p_(corrected)_ = 0.007). There was no significant difference in neural entrainment to the spectral dimension between attend pitch and neutral conditions (ITPC: V = 141, z = 0.886, p_(corrected)_ = 0.428, amplitude: V = 123, z = 0.261, p_(corrected)_ = 0.812).

Thus, for both pitch and spectral peak dimensions, attention enhanced neural entrainment to the attended dimension. However, there was no evidence that the unattended dimension was suppressed relative to the neutral attention condition.

### Correlational analyses

#### Relationship between Behavioral and EEG Data

To determine whether there was a relationship between behavioral performance and neural entrainment, we tested for a correlation between d-prime and the size of the attention effect in attend pitch and spectral peak conditions. Spearman’s correlations were used with FDR correction for multiple correlations (Benjamini & Hochberg, 1995). In the attend pitch condition, the neural attention effect was measured by taking the difference in pitch entrainment between conditions in which pitch was attended (attend pitch) and unattended (collapsed across attend spectral peak and neutral conditions). In the attend spectral peak condition, the neural attention effect was measured by taking the difference in spectral peak entrainment between conditions in which spectral peak was attended (attend spectral peak) and unattended (collapsed across attend pitch and neutral conditions). Correlations were conducted for both ITPC and amplitude data.

These analyses did not show any significant relationship between d-prime and neural entrainment in either the attend pitch condition (ITPC: rho = 0.484, p_(corrected)_ = 0.104; amplitude: rho = 0.392, p_(corrected)_ = 0.157) or attend spectral peak condition (ITPC: rho = − 0.136, p_(corrected)_ = 0.654; amplitude: rho = 0.104, p_(corrected)_ = 0.654).

#### Relationship with musical training

In this experiment, we predominantly recruited participants who received musical training. This provided the opportunity to examine whether there was a relationship between dimension-selective attention and years of musical training.

As shown in Figure 5, there was a significant positive correlation between musical training and d-prime in the attend pitch condition (rho = 0.55, p_(corrected)_ = 0.010) and attend spectral peak condition (rho = 0.53, p_(corrected)_ = 0.013).

**Figure 5.**
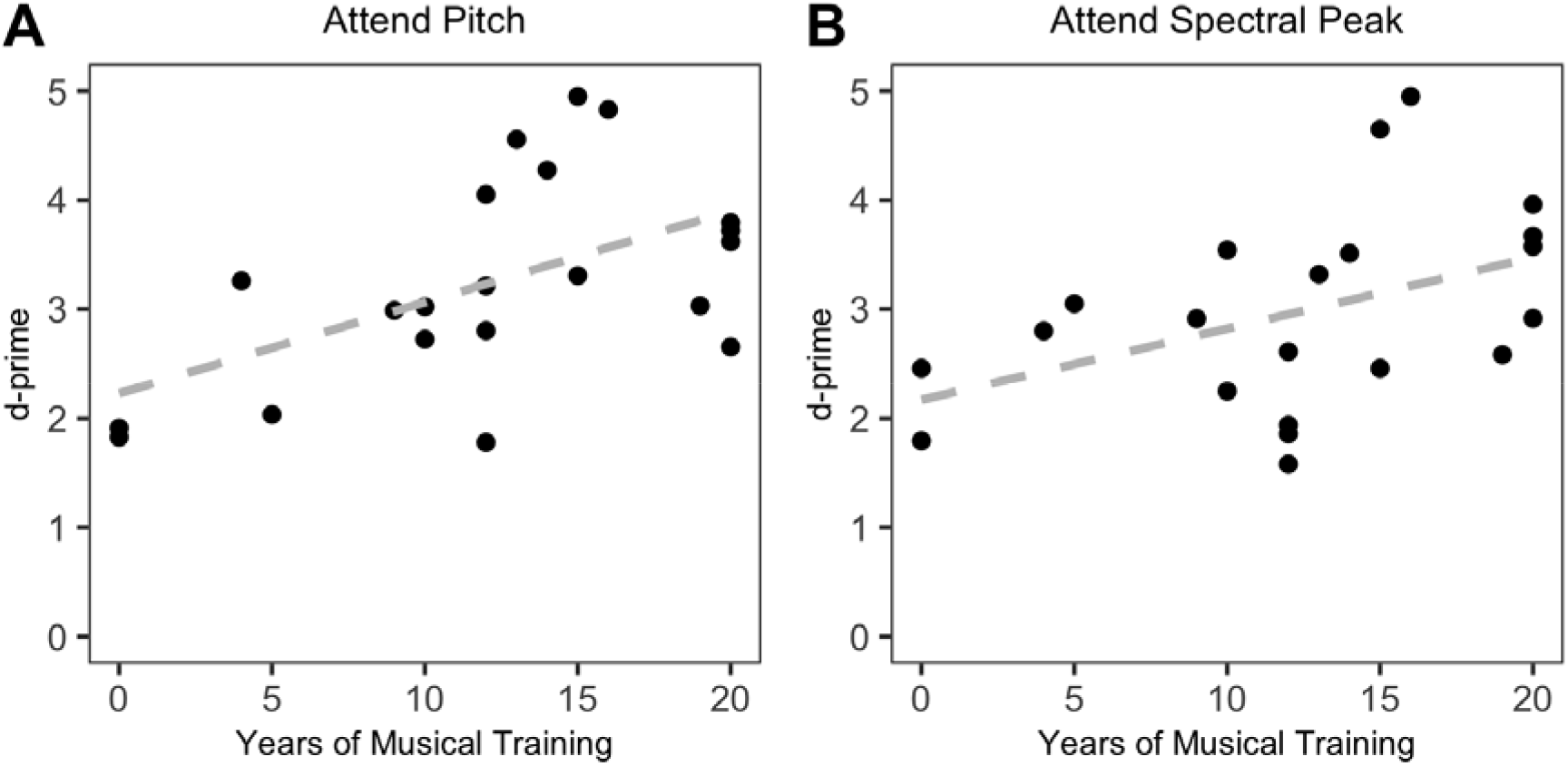
Relationship between years of musical training and d-prime on the dimension-selective attention task in attend pitch (A) and attend spectral peak (B) conditions. The correlation between years of musical training and d-prime was significant in the attend pitch condition, suggesting that individuals with more musical training were better at selectively attending to both acoustic dimensions.

To explore whether this relationship with musical training was observed in the neural data, we correlated years of musical training with the attention effect in attend pitch and attend spectral peak conditions. However, there was no significant correlation between years of musical training and the attention effect for pitch (ITPC: rho = 0.432, p_(corrected)_ = 0.125; amplitude: rho = 0.413, p_(corrected)_ = 0.125) or spectral peak (ITPC: rho = −0.044, p_(corrected)_ = 0.848; amplitude: rho = −0.099, p_(corrected)_ = 0.848) after correcting for multiple correlations.

### Discussion

In two studies, we investigated the neural correlates of dimensional salience and selective attention within a single sound stream that varied in pitch and spectral peak frequency. Consistent with our hypotheses, both dimensional salience and dimension-selective attention modulated neural entrainment. Specifically, Experiment 1 showed stronger neural entrainment to changes along the pitch dimension for tone sequences with high pitch salience (2-semitone pitch step sizes) compared to low pitch salience (1-semitone pitch step sizes). By contrast, relative pitch salience did not affect entrainment to simultaneously occurring changes in a different dimension, spectral peak frequency.

In Experiment 2, we found stronger entrainment for attended compared to unattended dimensions. These findings are consistent with previous studies showing that both bottom-up and top-down attentional mechanisms modulate neural entrainment to distinct auditory objects (Elhilali et al., 2005; Huang & Elhilali, 2020; Kaya et al., 2020). Here we extend these findings to show that dimensional salience and dimension-selective attention can enhance neural entrainment to different acoustic dimensions within a single sound stream.

There are many potential factors that may drive dimensional salience. Here we focus on pitch, which is one among the most important acoustic dimensions for human language and music (Patel, 2010). Previous research has shown that deviations in pitch can capture attention to auditory streams (Berti et al., 2004; Escera et al., 1998; Marois et al., 2018; Siddle et al., 1984), with the size of pupil dilation responses (Marois et al., 2018; Wetzel et al., 2016) and ERP responses (Berti et al., 2004; Rinne et al., 2006; Schröger, 1996) varying in proportion to pitch step size. Moreover, a recent EEG study has observed an increase in power and cross-trial coherence in response to deviant tones with high (+6 semitones) compared to low (+2 semitone) pitch changes (Kaya et al., 2020). Our results are consistent with these findings, showing that changes in the pitch step size elicit stronger neural entrainment specifically to pitch variations. An advantage of our paradigm is that it does not require deviant changes in an acoustic dimension to measure dimensional salience, but rather measures salience across entire recording blocks. To further establish this increase in neural entrainment as a marker of dimensional salience, future studies could present acoustically identical stimuli while altering the perceived salience of different acoustic dimensions within or between-subjects. For instance, this paradigm could provide a measure of dimensional salience before and after learning novel non-speech auditory categories that vary in the reliability of different acoustic dimensions (Holt & Lotto, 2006), or at different stages of second language acquisition. Alternatively, this measure could be used to explore differences in dimensional salience that occur between speakers of different languages, such as pitch contour for speakers of tone languages versus non-tone languages.

In addition to being modulated by dimensional salience, neural entrainment was also enhanced by dimension-selective attention. This effect of attention was demonstrated for two distinct dimensions – pitch and spectral peak. Previous research has shown that pitch and spectral dimensions can interact, with random variations in one dimension impeding discrimination of the other (Allen & Oxenham, 2014). However, our results provide one of the first demonstrations that attention can be directed to different acoustic dimensions. Whereas previous studies of auditory selective attention have established neural correlates of auditory stream segregation (e.g., Ding et al., 2016; Elhilali et al., 2005; Kerlin et al., 2010; Laffere et al., 2020, 2021; Zion Golumbic et al., 2013), here we show reliable neural correlates of attention to acoustic dimensions within a single sound stream.

The inclusion of the neutral attention condition was aimed at testing the extent to which the effects of attention were driven by neural enhancement of the attended dimension versus suppression of the unattended dimension (Chait et al., 2010). Previous research of auditory object-based attention (Horton et al., 2013) has observed suppression effects of neural entrainment to unattended stimuli. Suppression has also been suggested to play a role in attention to different acoustic dimensions (Costa-Faidella et al., 2017). However, we observed no evidence of suppression. This suggests that, at least in the context of the current study, the neural entrainment effects were driven by enhanced gain of the attended dimension rather than active suppression of the unattended dimension.

While our results show that attention can be directed to pitch and spectral peak, it remains to be seen whether these results will generalize to other acoustic dimensions. Futures work can examine the extent to which the effects of dimensional salience and dimension-selective attention generalize to other acoustic dimensions. Another potential limitation to the current study stems from the difficulty of the dimension-selective attention task. Behavioral results show that participants can selectively attend to pitch and spectral peak dimensions, with performance at above chance levels. However, dimension-selective attention appears to be a difficult task for many individuals. Of our initial sample of 27 participants, 6 failed to reach the required performance threshold on the training tasks. For this reason, we predominantly recruited individuals with musical training backgrounds. To explore this effect in a wider sample of participants, future studies could increase the amount of training provided and/or decrease the difficulty of the task by slowing the rate of tone presentation and decreasing the number of levels of each dimension.

The variability in dimension-selective attention performance across individuals raises the question of whether and how dimension-selective attention can be trained. Previous research has shown that relatively short-term training can improve speech perception in noise (Whitton et al., 2014, 2017) and enhance electrophysiological indices of auditory spatial attention (Isbell et al., 2017; Stevens et al., 2008, 2008). Even two hours of training can boost behavioral and neural measures of auditory selective attention (Laffere et al., 2020). If training can improve object-based auditory attention, then it may be possible to train dimension-selective attention as well. Indirect support for this idea comes from studies showing that categorization training can alter the weight listeners place on different acoustic cues (Chandrasekaran et al., 2010; Francis et al., 2000, 2008). The paradigm used in the current study could provide a more direct measure of training-induced changes in dimension-selective attention. If training can boost dimension-selective attention, it could be used to help improve second-language learning by facilitating attention to acoustic dimensions relevant for categorization.

We also observed large individual differences in neural measures of dimensional salience and dimension-selective attention, consistent with previous studies showing large individual differences in object-based attention (Choi et al., 2014; Laffere et al., 2020; Ruggles & Shinn-Cunningham, 2011; Tierney et al., 2020). There are multiple potential factors that may contribute to these individual differences. On the one hand, individual differences in cortical tracking of a particular dimension may reflect variability in the precision with which that dimension is encoded by the auditory system. In this case, we would predict individuals who have more difficulty encoding a particular dimension (e.g., individuals with congenital amusia who have difficulty processing pitch) to show less passive cortical tracking of that dimension and potentially be less sensitive to variations in the salience of that dimension. On the other hand, variability in executive function or attentional control may contribute to differences in cortical tracking of attended dimensions. Since there is little research explicitly investigating dimension-selective attention, it is unclear whether and how this type of attention relates to other forms of auditory attention (e.g., object-based attention) as well as other executive functions. Understanding how these sensory and cognitive factors contribute to individual differences in dimensional salience and dimension-selective attention, and in turn, how these individual differences relate to auditory perception in natural listening environments remains a key topic for future research.

The study of dimensional salience and dimension-selective attention has implications for speech perception and language acquisition. Attentional accounts of cue weighting suggest that listeners selectively attend to acoustic dimensions that are most diagnostic of category identity (Francis et al., 2000; Francis & Nusbaum, 2002; Holt et al., 2018). However, research investigating the link between dimension-selective attention and cue weighting in speech perception is sparse, in part because of a lack of established measures of dimension-selective attention. The paradigms used in our studies could be coupled with studies of cue weighting in speech perception to establish an association between dimensional salience, selective attention and cue weighting. For example, a study could train participants to learn novel categories that differ along two (or more) dimensions and examine whether cue weights correlate with phase-locking to each dimension. Alternatively, research using speech stimuli could examine whether short-term changes in cue distribution result in related changes in neural entrainment.

This paradigm could also be used to compare dimension-selective attention ability across different populations, such as individuals with and without musical training. In Experiment 2, we predominantly recruited participants who received musical training because of the difficulty of the task. This allowed us to test whether there was a relationship between years of musical training and dimension-selective attention. Musical training was associated with improved behavioral dimension-selective attention performance for both pitch and spectral peak dimensions. That this effect was not observed in the neural data may have been due to lack of power, as this study was not explicitly designed to test for an effect of musicianship. Nonetheless, the behavioral effect suggests that musical training may be associated with robust dimension-selective attention. Evidence from previous research suggests that musical training is associated with less informational masking during tone detection (Oxenham et al., 2003), improved nonverbal auditory selective attention (Tierney et al., 2020), and better speech-in-speech perception in some studies (Clayton et al., 2016; Du & Zatorre, 2017; Parbery-Clark et al., 2009; Slater & Kraus, 2016; Zendel & Alain, 2012; but see Boebinger et al., 2015; Fuller et al., 2014; Madsen et al., 2017; Ruggles et al., 2014). Our results provide an initial indication that this musician advantage might extend to dimension-selective attention. This paradigm could be used to further explore the relationship between musical training and dimension-selective attention. For example, future studies could compare trained musicians to non-musicians, or potentially compare entrainment to variations in pitch salience in musicians who specialize in melodic instruments versus percussionists.

In conclusion, we provide evidence that dimensional salience and dimension-selective attention modulate neural entrainment, with stronger entrainment to salient and attended dimensions. This study offers a paradigm that can be used to explore the effects of dimensional salience and dimension-selective attention across multiple acoustic domains.

## Acknowledgements

This work was supported by a Research Project Grant from the Leverhulme Trust [grant number RPG-2019-107] to ATT and a Reg and Molly Buck Award from SEMPRE to AES.

## Footnotes

1 One participant who failed to reach the performance threshold in the single dimension training but went onto reach the threshold in the dimension-selective attention training was included in this experiment.

