## Supplementary Materials for "Dimension-Selective Attention and Dimensional Salience Modulate Cortical Tracking of Acoustic Dimensions"

### S.1. Differences between online and in-lab training protocols

#### Overview

The online training protocol was designed to mirror the in-lab protocol as closely as possible. However, some minor changes needed to be made due to the change in platforms. Most notably, the in-lab training was presented using PsychoPy3 while the online training was presented using Gorilla. This required us to switch from using .wav files to .mp3 files, since .wav files are not supported on the Gorilla platform. Moreover, in the lab, participants used ear inserts for the training as well as EEG recording. Since the online training was completed at the participants’ homes, each participant used their own headphones to listen to the stimuli. Finally, responses were made on an Xbox game controller in the in-lab training but on a keyboard in the online training. Further details of the design of each training task are provided below.

#### In-lab Training

In both the single dimension and dimension-selective attention training exercises, participants listened to 48-second sequences of tones. For attend pitch and spectral peak conditions, participants were instructed which dimension to attend to and how often that dimension would change. Their task was to press the trigger button on an Xbox gamepad when they detected repetitions in the attended dimension. In the neutral condition, participants were instructed to listen out for occasional quiet tones and press a button when they detected a quiet tone.

Participants received visual feedback on their performance. When participants correctly identified a repetition/quiet tone, the word ‘Hit!’ was displayed on the screen. When participants missed a repetition/quiet sound, the word ‘Miss!’ was displayed on the screen. Participants’ score (number of hits and false alarms) was displayed in the right-hand corner of the screen. Participants received +1 for each it and -1 for each false alarm to provide a total score.

For the attend pitch and attend spectral conditions, the two rates of change (1.33 and 2 Hz) were presented in alternating manner. If participants failed to reach the performance threshold (performance collapsed across the two rates of dimension change), they received another training block for that condition. Once they reached the threshold, they moved onto the next condition. If participants did not pass all of the single-dimension training conditions, they did not progress to the dimension-selective attention training. If participants did not pass all of the dimension-selective attention training conditions, they were reimbursed for their time but did not take part in the EEG task.

#### Online Training

The stimuli were identical to the stimuli presented in the in-lab training, with the exception that the audio files were converted to mp3 format. The task was also the same. In the attend pitch and spectral peak conditions, participants used their mouse to press a button on the screen when they detected repetitions in the attended dimension. In the neutral condition, participants were instructed to listen out for occasional quiet tones and press the button on the screen when they detected a quiet tone.

Participants received visual feedback on their performance. When participants correctly identified a repetition/quiet sound, the word ‘Good!’ was displayed on the screen. When participants clicked the button when there was not a repetition/quiet sound, the word ‘Oops!’ was displayed on the screen. Participants’ score for that block and their total score for that condition was displayed on the screen.

For both training tasks, stimulus presentation was blocked by condition (attended dimension and rate of change). If participants failed to reach the performance threshold (performance not collapsed across the two rates of dimension change), they received another training block for that condition. Once they reached the threshold, they moved onto the next condition. If participants did not pass all of the single-dimension training conditions, they did not progress to the dimension-selective attention training. If participants did not pass all of the dimension-selective attention training conditions, they were reimbursed for their time but did not take part in the EEG task.

#### Results

We did not expect the subtle differences in training procedure to influence training performance. To ensure this was the case, we examined the hit rate for online versus in-lab training collapsed across attend pitch and spectral peak conditions. For the single dimension training, the hit rate was comparable for online (median = 96.9% mean = 95.7%, sd = 6.2%) and in-lab (median = 96.9%, mean = 97.2%, sd = 3.3%) participants. Similarly, performance on the dimension-selective attention training was comparable for online (median = 90.6%, mean = 90.9%, sd = 4.5%) and in-lab (median = 96.9%, mean = 94.5%, sd = 5.7%) participants.

The difference between online and in-lab training that was most likely to influence performance was the fact that in-lab training was the temporal delay between training and EEG recording. While in-lab participants completed training immediately prior to EEG recording, online participants had a delay between training and EEG recording. This could have resulted in a deterioration in performance between training and testing for the online participants. However, this did not appear to be the case. Dimension-selective attention performance of the participants who completed the training online (n = 7, median = 82.5%, mean = 83.5%, sd = 10.8%) was similar to the performance of the participants who completed the training in-lab (n = 14, median = 82.5%, mean = 83.2%, sd = 12.7%). Thus, the delays between training and testing in online participants did not appear to adversely affect task performance.

Figure S.1. shows the phase-locking data for participants in each training group. Although it is difficult to compare the two groups given that there are twice as many in-lab as online training participants, the pattern of results appear broadly consistent irrespective of training.

These results suggest that the subtle differences in training protocols required due to the COVID-19 pandemic did not substantially affect behavioral or neural data.

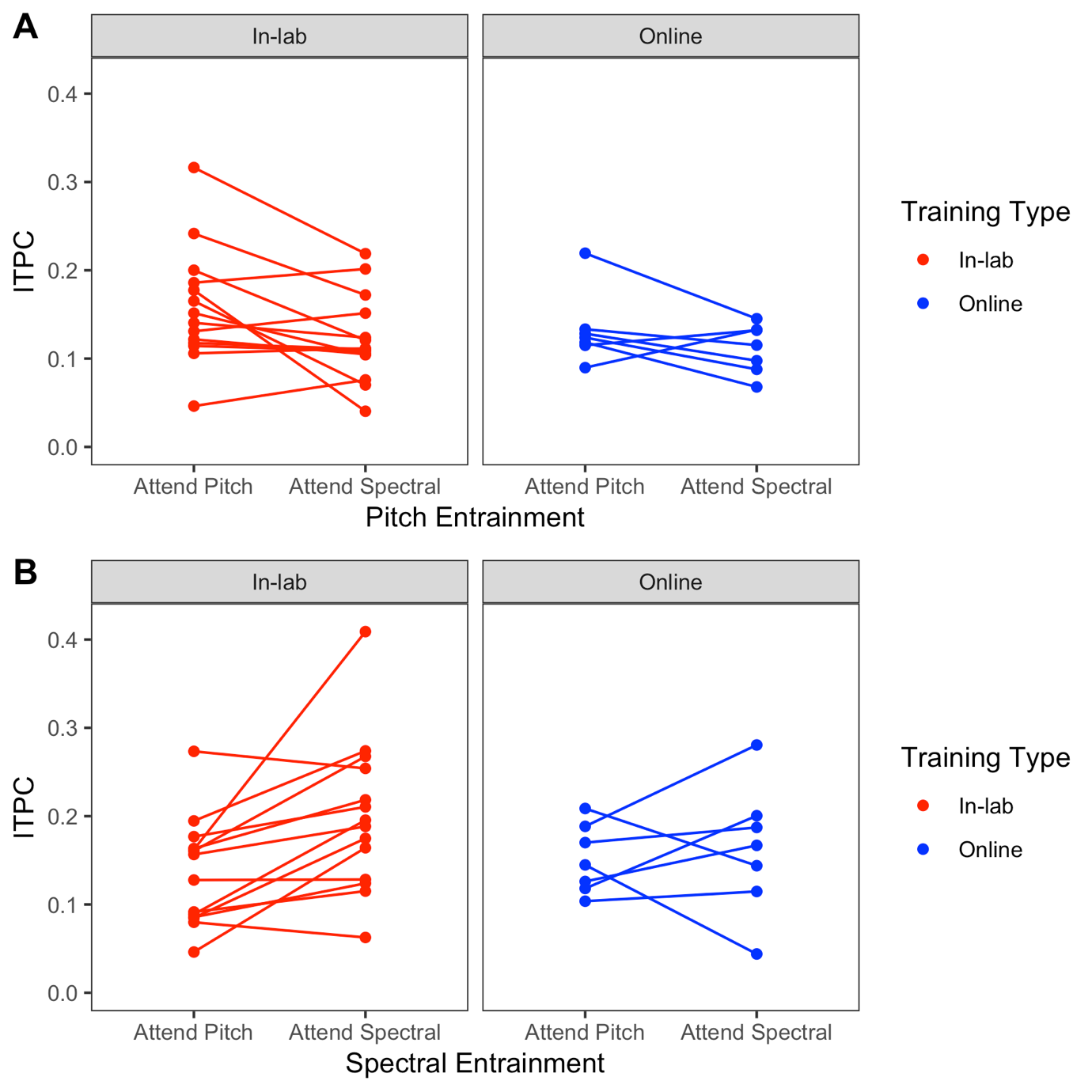

Figure S.1. Neural entrainment (ITPC) to pitch (A) and spectral peak (B) dimensions in the attend pitch and spectral peak conditions (x-axis). The lines show individual participants who received online versus in-lab training.
